# Neuromodulator control of energy reserves in dopaminergic neurons

**DOI:** 10.1101/2025.08.19.671144

**Authors:** Camila Pulido, Matthew S. Gentry, Timothy A. Ryan

## Abstract

The brain is a metabolically vulnerable organ as neurons have both high resting metabolic rates and the need for local rapid conversion of carbon sources to ATP during activity. Midbrain dopamine neurons are thought to be particularly vulnerable to metabolic perturbations, as a subset of these are the first to undergo degeneration in Parkinson’s disease (PD), a neurodegenerative disorder long suspected to be in part driven by deficits in mid-brain bioenergetics (1). In skeletal muscle, energy homeostasis under varying demands is achieved in part by its ability to rely on glycogen as a fuel store, whose conversion to ATP is under hormonal regulatory control. In neurons however the absence of easily observable glycogen granules has cast doubt on whether this fuel store is operational, even though brain neurons express the key regulatory enzymes associated with building or burning glycogen (2). We show here that that in primary mid brain dopaminergic neurons, glycogen availability is under the control of dopamine auto receptors (D2R), such that dopamine itself provides a signal to store glycogen. We find that when glycogen stores are present, they provide remarkable resilience to dopamine nerve terminal function under extreme hypometabolic conditions, but loss of this dopamine derived signal, or impairment of access to glycogen, makes them hypersensitive to fuel deprivation. These data show that neurons can use an extracellular cue to regulate local metabolism and suggest that loss of dopamine secretion might make dopamine neurons particularly subject to neurodegeneration driven by metabolic stress.

**Significance Statement:** This work demonstrates that a neuron’s metabolic resilience is actively shaped by local neuromodulation, providing a potential explanation for why different neuronal subtypes exhibit unequal vulnerability to metabolic stressors and why dopamine neurons become more vulnerable when they lose the capacity to release dopamine.

## Introduction

Glycogen, an osmotically neutral polymer of glucose, is the major glucose reserve in eukaryotes, and defects in glycogen metabolism and structure lead to disease. The fundamental understanding of glycogen metabolism stems from pioneering studies in liver and muscle over the last 80 years (3). In liver, glycogen serves as the fuel store to maintain blood glucose homeostasis, where insulin and glucagon mediate the balance of glycogen synthesis or breakdown resulting in net glucose uptake or export from this organ. Muscle uses glycogen as a fuel store that can rapidly mobilize phosphorylated glucose from the glucose polymer under the control of epinephrine (4). The liberated glucose is then oxidized to ATP and pyruvate to meet the bioenergetic needs of muscle contraction. Neurons, like muscle, have bioenergetic needs that vary over time and must synthesize ATP on demand to meet the needs of the neuronal molecular machinery engaged by electrical activity (5). Failure to meet the local ATP needs in axons during activity leads to rapid collapse of synaptic function (6, 7). Paradoxically, the brain, and in particular neurons, have long been considered to rely minimally on glycogen stores despite expressing the key enzymes associated with glycogen metabolism (2). Part of this view stems from the fact that the classic way to visualize glycogen is by electron microscopy. In resting muscle for example, glycogen polymers of up to 50,000 glucose molecules are synthesized and can be easily recognized as a classic granule signature (8), but smaller granules are not easily distinguished from other electron-dense subcellular material. Although glycogen granules are occasionally visualized in astrocytes, particularly under certain forms of anesthesia (9), they are not readily observed in neurons, leading to the prevailing view that most glycogen metabolism in the brain resides exclusively in non-neuronal cells. It is important to note however, that the abundance of glycogen reflects the steady-state balance of synthesis and breakdown, and low steady-state abundance may simply reflect a constant flux through the pathway. The consequence of disrupting glycogen metabolism in the brain on neuronal function has usually been interpreted as evidence for the shuttling of a glycolytic end-product, lactate from astrocytes (10). Recent quantitative evidence however does not support the idea that astrocytes, as a rule, provide lactate during or after electrical activity to support neuronal function (11, 12). Additionally, neuron-specific genetic ablation of critical proteins needed for glycogen metabolism leads to significant impairment in cognitive function (13) supporting the notion that neuronal glycogen metabolism plays an important role in normal brain function. Although glycogen metabolism would be expected to lead to increased ATP production, as is likely in muscle treated with epinephrine, recent evidence indicates that under some circumstances, glycogen metabolism is shunted to the pentose phosphate pathway, in turn providing protection against oxidative stress (14, 15). To help clarify whether or not neuronal glycogen metabolism plays a role similar to that in muscle, we made use of primary neurons, where it is possible to acutely drive bioenergetic needs with electrical activity while monitoring intracellular ATP dynamics (16) or synaptic function (17). Here we show that primary rat mid-brain dopamine neurons can rely on glycogen to sustain nerve terminal function in the absence of glucose, but that the ability to do so depends on the previous activity of D2Rs. Pretreatment of these neurons with a D2R antagonist, leads to depletion of glycogen stores, and makes their nerve terminals hypersusceptible to loss of extracellular glucose, and unable to sustain cytoplasmic ATP production. These data provide direct evidence that glycogen, even in the absence of external glucose, can meet the bioenergetic needs of synapses under prolonged stimulation, but access to glycogen in turn is under neuromodulatory control. The data suggest that as dopamine secretion becomes impaired in the brain, the loss of the autocrine signal in turn will make dopamine neurons more susceptible to metabolic stressors. The critical importance of D2Rs in regulating this energy reserves additionally suggests this may underlie the susceptibility of presymptomatic patients to drug-induced Parkinsonism driven by atypical antipsychotics that target D2Rs (18, 19).

## Results

### Primary Dopaminergic axon terminals are highly resistant to glucose deprivation

As mid-brain dopaminergic neurons are known to undergo preferential degeneration in PD, and PD is thought in part to be driven by deficits in brain bioenergetics, we sought to examine the vulnerability of dopamine neurons to specific metabolic stresses using a reductionist approach *in vitro*. We previously developed a quantitative optical approach to examine the sensitivity of synaptic performance to metabolic perturbations, using the dependence of vesicle recycling on ATP availability (17, 20). Here we made use of the tyrosine hydroxylase (TH) promoter (21) to specifically enrich for expression of different genetically encoded functional indicators in dissociated mid-brain rat dopamine neurons (Fig. S1) to examine their sensitivity to metabolic stress, in particular to hypoglycemic conditions. We examined the kinetics of synaptic vesicle (SV) recycling during repeated bouts of stimulation (50 APs, 10 Hz) delivered at minute intervals in the absence of external glucose. Consistent with our previously work (20), this protocol leads to an arrest in synaptic vesicle retrieval in hippocampal neurons after ∼5 rounds of stimulation (Fig. 1A). To characterize the variation in synaptic endurance across cells, we measured the fraction of the exocytic signal that remained after a defined post stimulus time period (3 × the decay time constant in one round of stimulation in 5mM glucose, 3τ), which we term the percentage of endocytic block (EB) (= ∼5% in the first round by definition for a perfect exponential decay). For a given neuron, we determined the number of stimulus rounds it took for SV recycling to fail to retrieve 50% of the exocytic signal at this time period (EB in the trace), which in hippocampal neurons was, on average, 5.95 rounds ± 0.45 (n = 20; Fig. 1C) consistent with previous findings (20). In contrast we were surprised to discover that dopaminergic nerve terminals exhibited a much greater resilience to the withdrawal of glucose and sustained robust SV recycling on for many rounds of repeated burst of AP firing, where a large subset (8/29) failed to arrest for all 30 rounds of stimulation (Fig. 1 B, C).

**Figure 1.**
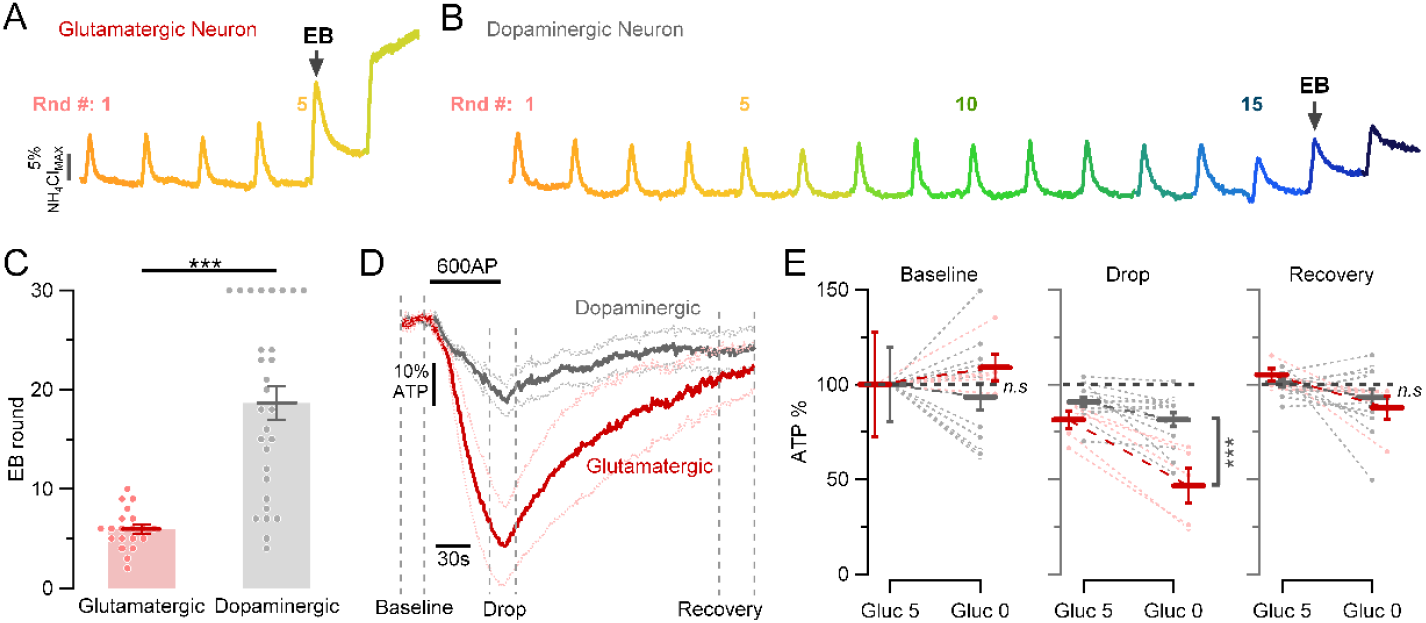
Dopaminergic axonal terminals are more resilient to fuel deprivation compared to Glutamatergic terminals. **(A-B)** Synaptic function measured in neurons expressing Syphy-pH in the absence of glucose (0mM glucose). Color-coded traces represent responses to one stimulus round (50 Aps, 10 Hz) applied every minute (from orange = first round to dark blue = seventeenth round). Responses are normalized to sensor expression using ammonium chloride measurements. (A) Trace to six stimulus rounds in a glutamatergic neuron. (B) Trace to seventeen stimulus rounds in a dopaminergic neuron. (C) Number of rounds of stimulation before endocytic block exceeds 50% is smaller in glutamatergic neurons (red dots, n = 20) than dopaminergic neurons (gray dots, n = 29): mean ± SEM: 5.95 ± 0.45 rounds versus 18.66 ± 1.69 rounds, where 8 out 29 dopaminergic neurons never arrested for all 30 rounds of stimulation. (D) Average axonal ATP changes to 600 APs, 10Hz stimulation in neurons expressing iATPSnFR2.0 in the absence of glucose (0mM glucose), normalized to pre-stimulation baseline. Dopaminergic (gray trace, n = 14) versus glutamatergic axonal terminals (red trace, n = 5). (E) ATP dynamics before (Baseline), during (Drop) and after (Recovery) electrical activity in 5mM glucose (Gluc 5) and after 5min incubation without glucose (Gluc 0). Dopaminergic axons generate sufficient ATP during and after electrical activity to sustain SV recycling than glutamatergic axons. ATP drop during stimulation in absence of glucose in dopaminergic vs Glutamatergic: mean ± SEM: 18.64 ± 3.69 vs 53.4 ± 9.16 respectively. ^***^P<0.001, non-significant as ns, Wilcoxon-Mann-Whitney test.

The metabolic resilience of the primary dopamine neurons strongly implies that even in the absence of external glucose, their nerve terminals generate sufficient ATP during and after electrical activity to sustain SV recycling. To test this hypothesis, we expressed a recently developed genetically encoded fluorescent ATP sensor, iATPSnFR2.0 (16) in dopamine neurons. As shown previously, when glutamatergic hippocampal neurons are subjected to prolonged stimulation in the absence of external glucose, ATP levels drop precipitously (∼53 %) and recover only slowly after the cessation of activity, consistent with the idea that there is no readily available oxidizable carbon source to synthesize ATP as it is consumed (Fig. 1D, E). In contrast, in dopamine neurons, ATP levels decrease by only ∼18% using the same stimulus paradigm and readily returned to baseline following the electrical activity (Fig. 1 D, E). These data thus demonstrate that primary mid-brain dopamine neurons deploy a mechanism that allows them to generate ATP during activity in the complete absence of an explicit external carbon source that is sufficient to sustain robust nerve terminal function.

### Dopamine neurons and their nerve terminals show robust expression of glycogen synthase and glycogen

The remarkable metabolic resilience of dopamine neurons even when bathed in glucose-free saline suggests that these neurons have access to an intracellular source of combustible carbon to maintain synaptic function. We tested the idea that glycogen might be the relevant carbon source. Although electron microscopy is the traditional approach to detect the classic granule signature, we reasoned that this might be difficult to implement for *in vitro* studies, as the primary dissociated dopamine neurons are part of a mixed cell type, that include both astrocytes and non-TH expressing neurons. Additionally, when glycogen granules are only a few nanometers in size they are indistinguishable from other electron-dense material in the cytoplasm. To overcome this problem, we utilized a monoclonal antibody previously shown to have very high affinity for both purified glycogen granules as well as enzymatically digested glycogen (22, 23). We triple-labeled primary mid-brain dissociated neuron cultures with the anti-glycogen, anti-TH and an antibody for glycogen synthase (GS). These experiments showed directly that dopamine neurons (TH-positive) have robust expression of GS and have an accumulation of glycogen (Fig. 2A). Glycogen and GS were distributed through the somato-dendritic regions and in axons, including at presynaptic varicosities. Both the amount of glycogen and glycogen synthase detectable in individual cell somas from a mix population (TH positive and negative) was quite variable with more than an order of magnitude difference in each of these across cells (Fig. 2B). Analysis of the paired intensities of glycogen and GS over >130 individual cells showed that there is a linear correlation between accumulation of glycogen and the expression of GS (Fig. 2B). We hypothesized that dopamine neurons were particularly enriched in glycogen. To examine this, we compared the intensities of glycogen staining on the same coverslip on neurons that were TH positive and negative (Fig. 2C). This analysis showed that on average dopamine neurons have ∼ 70% more glycogen expression than non-TH-positive neurons (Fig. 2D).

**Figure 2.**
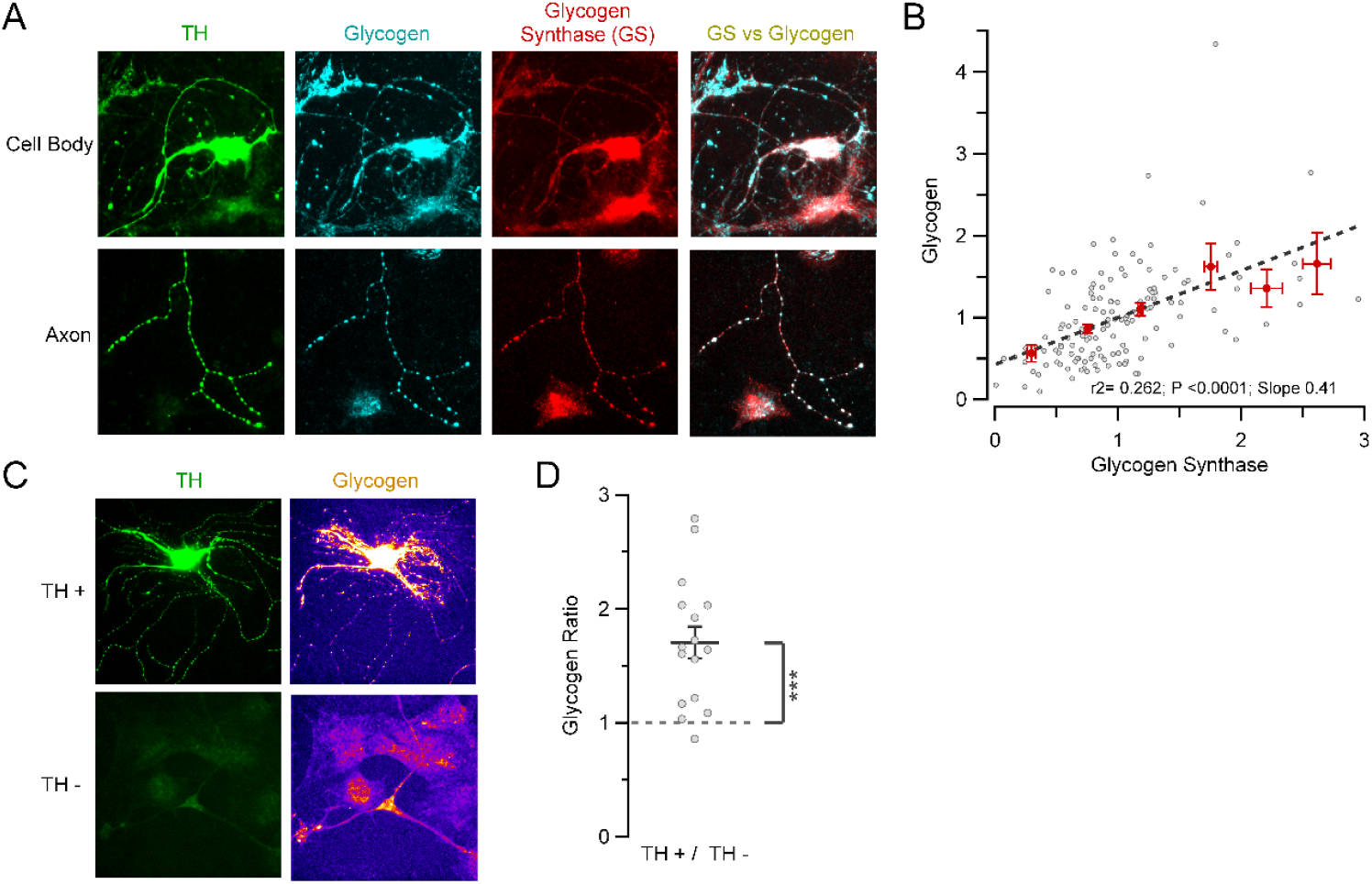
Glycogen synthase is expressed in dopaminergic axon terminals, enabling local glycogen storage. (A) Representative cell body (top) and axon (bottom) images tripe-labeled with anti-TH (green), anti-Glycogen (cyan) and anti-Glycogen Synthase (GS; red) shows that dopaminergic terminals have the enzymatic machinery to synthase and store glycogen. Overlay image between GS vs glycogen is shown in the right column. (B) Glycogen storage levels in cell bodies from a mix of TH+ and TH-neurons (gray dots; n = 138 cell bodies from 5 cultures) are linearly correlated (black dash line) with GS expression levels. Binned average by a 0.5 window (red dots; Mean ± SEM). (C) Representative images of a Dopaminergic neuron (TH+) versus a non-Dopaminergic neuron (TH -; green) from a same coverslip and their respective glycogen level content (color-code from dark violet = low levels to yellow = high levels; same scale top and bottom). (D) On average, dopaminergic neurons (TH+) storage more glycogen than non-dopaminergic neurons (TH-). TH+ / TH-ratio per dish: mean ± SEM: 1.7 ± 0.142; n = 16 coverslips from 9 cultures.

### Primary dopamine neurons utilize their own stored glycogen to sustain function under hypoglycemic conditions

The balance of glycogen abundance in tissues is set in part by the activities of glycogen synthase and glycogen phosphorylase (GP), with GP being required to liberate phosphorylated glucose monomers from the glycogen granule. GP activity is controlled by phosphorylase kinase whose activity is allosterically controlled by a combination of Ca^2+^ ions and protein kinase A (24). Immunostaining for GP shows that, like GS, it is expressed throughout axons and somato-dendric regions of primary mid-brain neurons (Fig. 3A). In order to test if glycogenolysis is supporting SV recycling and ATP production we made use of both an shRNA that led to a 50% reduction of GP (Fig. S3) as well as a small molecule glycogen phosphorylase inhibitor (GPI) and examined their impact on SV recycling and ATP production in the absence of glucose during electrical activity. Unlike in untreated control dopamine neurons, acute application of GPI or loss of GP expression (GP KD), resulted in SV recycling in the absence of glucose to arrest within 2 (GPI) or 7 (GP KD) rounds of stimulation (Fig. 3 B, C). Consistent with these observations, measurements of cytosolic ATP using iATPSnFR2.0 showed prolonged electrical activity in dopamine neurons treated with GPI led to a ∼70% reduction in ATP that failed to recover, while stimulation in GP KD neurons ATP levels dropped by ∼ 40% and recovered only very slowly following stimulation (Fig. 3 D, E).

**Figure 3.**
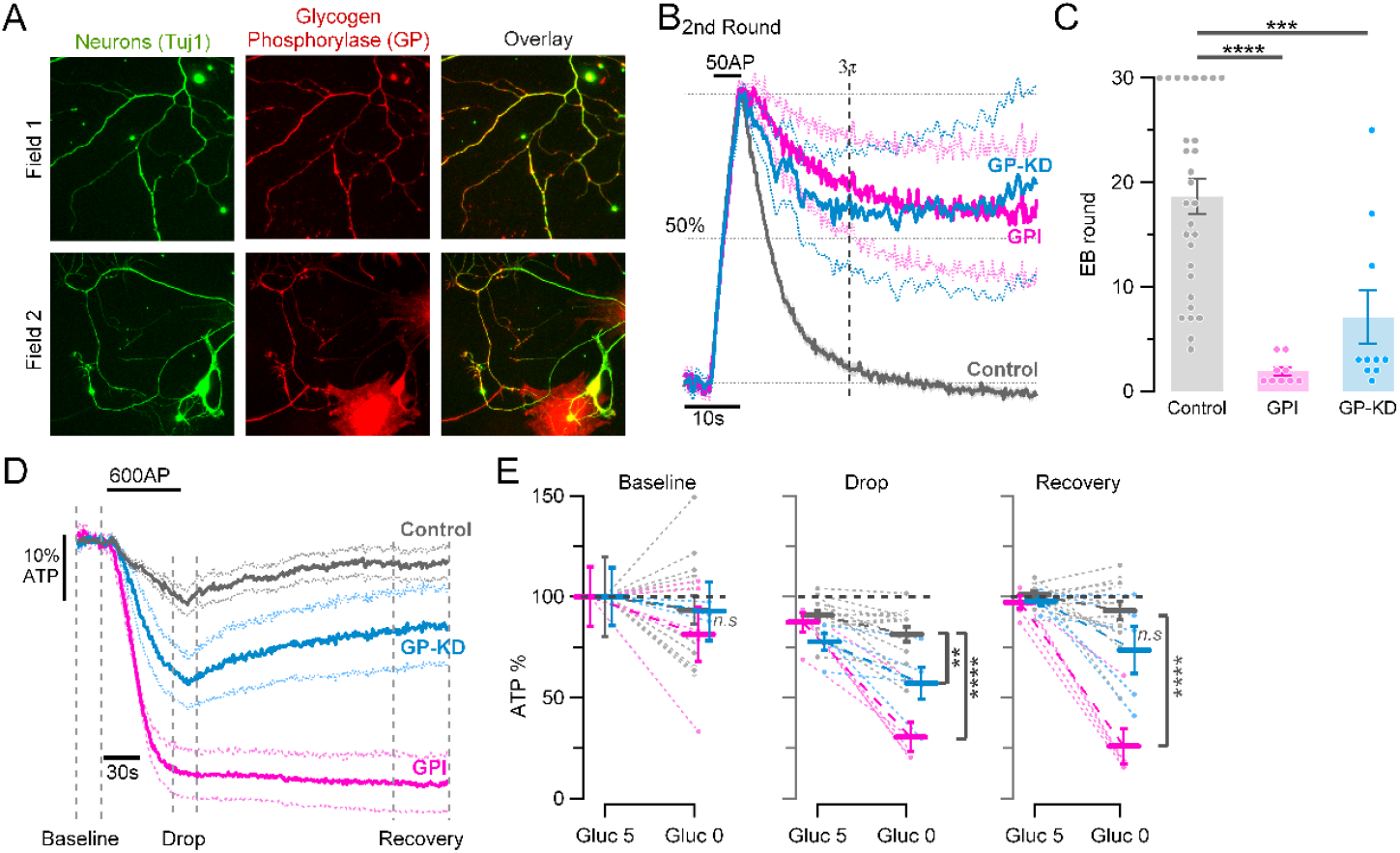
Dopaminergic terminals locally degrade glycogen into fuel to sustain synaptic function under hypoglycemic conditions. (A) Representative images of axons (Field 1 and 2) and somato-dendric regions (Field 2) labeled with anti-Tuj1 (green) and anti-Glycogen Phosphorylase (GP; red) shows that axonal terminals locally express the enzymatic machinery capable of breaking down glycogen into a fuel source. Overlay images are shown on the right column. (B) Average traces in absence of glucose at the second round of stimulation (50AP, 10Hz) in control neurons (gray trace; n = 29), neurons treated with GPI (magenta trace; n = 9) and neurons suppressing expression of GP (GP-KD; blue trace; n = 10). Responses are normalized to the peak. Vertical dashed line represents three times τ measured at 5mM glucose response (data not shown). Horizontal dashed line represents 50% retrieval of the exocytic signal. (C) Number of rounds of stimulation before endocytic block (EB) exceeds 50% is smaller in neurons with GP function impairment: GPI treatment and GP-KD (magenta and blue dots, respectively), than in neurons expressing functional GP (gray dots): mean ± SEM: 1.89 ± 0.42 rounds vs 7.1 ± 2.58 rounds vs 18.66 ± 1.69 rounds, respectively, where 8 out 29 dopaminergic neurons never arrested for all 30 rounds of stimulation. (D) Average axonal ATP changes to 600 APs, 10Hz stimulation in neurons expressing iATPSnFR2.0 in the absence of glucose, normalized to pre-stimulation baseline. Control (gray trace, n = 14) vs GPI treatment (magenta, n = 5) and GP-KD (blue, n = 5) neurons shows that dopaminergic axons depend on GP function to breakdown glycogen to be able to generate sufficient ATP to sustain activity. (E) Comparison of ATP percentage before (Baseline), during (Drop) and after (Recovery) electrical activity in 5mM glucose (Gluc 5) and after 5min in a glucose free saline solution (Gluc 0). ^****^P<0.0001, ^***^P<0.001, ^**^P<0.01, non-significant as ns, Wilcoxon-Mann-Whitney test.

### Glycogen storage in dopamine neurons is controlled by D2 auto-receptors

In muscle, the balance of glycogenesis versus glycogenolysis is controlled by the concentration of cAMP and Ca^2+^ ions as together they lead to the activation of GP. We reasoned that GPCRs that are coupled to Gi to inhibit cAMP production could potentially tilt the balance of activity towards glycogen storage. Mid-brain dopamine neurons express D2 receptors (25), a variant of dopamine receptor coupled to Gi/o (26). As dissociated primary dopamine neurons are likely exposed to a constant level of dopamine autocrine stimulation, perhaps similar to what occurs during tonic firing in the striatum (27), we reasoned that this stimulation of D2Rs might be sufficient to drive glycogen storage and form the basis of the metabolic resilience. To test this hypothesis, we made use of sulpiride, a clinically used antipsychotic D2R antagonist. Consistent with this idea, incubating primary dopamine neurons overnight in 1uM sulpiride led to a 50% reduction in glycogen content in axonal varicosities (Fig. 4A, B) and a ∼ 30% reduction in cell bodies (Fig. S4A, B). Notably, sulpiride had no impact on glycogen levels in non-dopaminergic neurons (Fig. S4B). To determine if this reduction in glycogen levels impacted axonal bioenergetics, we used our synaptic endurance test (Fig. 1 A, B) to examine the sensitivity of SV recycling to removal of glucose. These experiments showed that unlike with control neurons, previous treatment with sulpiride and the loss of glycogen, rendered primary dopamine neurons incapable of sustaining SV recycling for more than 2 rounds of stimulation (Fig. 4 C, D). Measurements of intracellular ATP revealed that neurons treated with sulpiride, even prior to stimulation, could not sustain ATP levels upon switching to a glucose free saline solution. During AP firing, the decline in ATP continued at an even faster rate, and failed to recover in the post-stimulus period in dopamine neurons treated with sulpiride (Fig. 4C E, F). These data strongly suggest that dopamine neurons, in the absence of access to glycogen, now are much more metabolically vulnerable. By comparison, hippocampal neurons do not change their metabolic sensitivity upon block with GP (Fig. S2).

**Figure 4.**
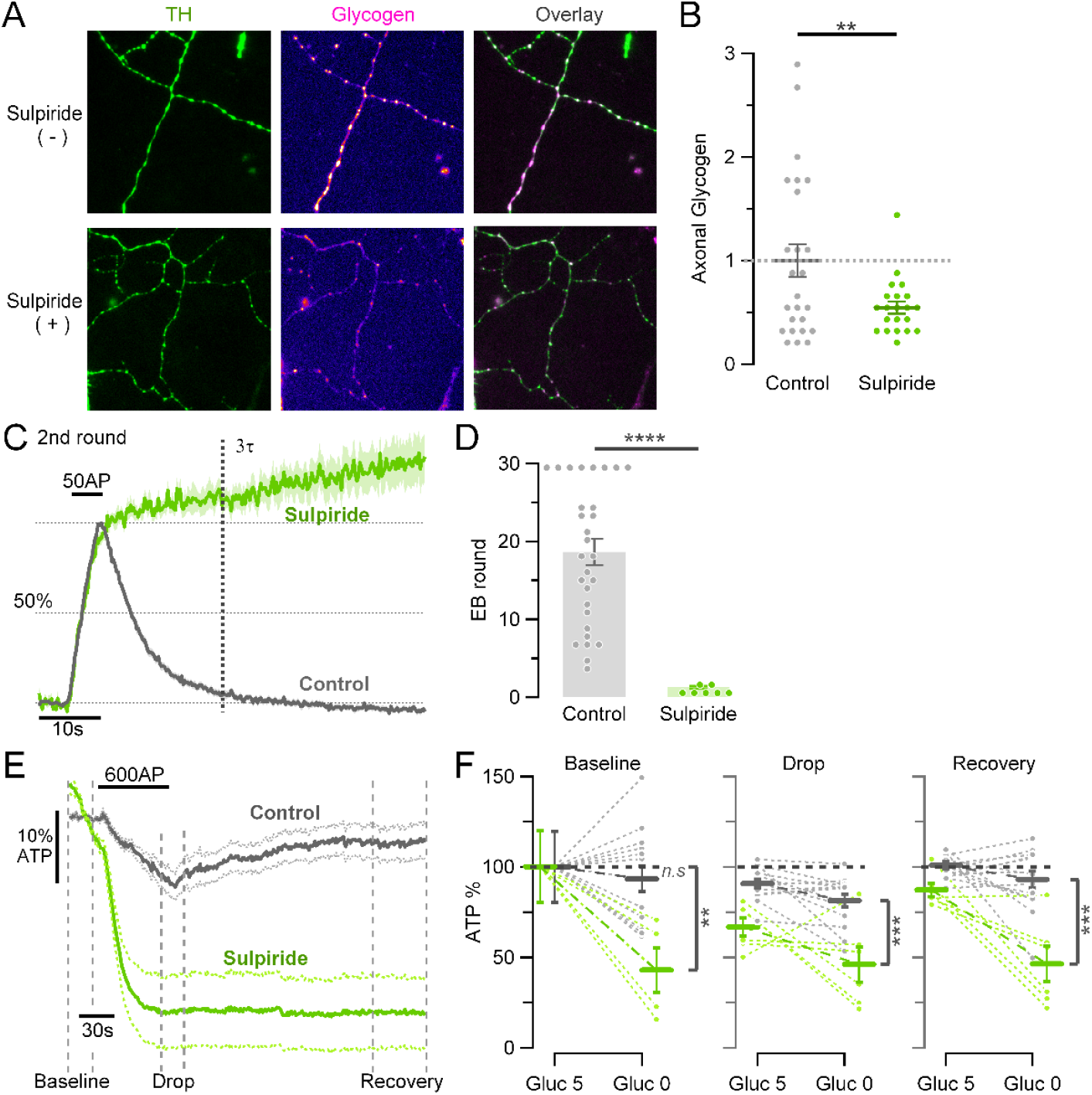
Local glycogen storage levels in dopamine neurons are modulated by D2 auto-receptors. (A) Representative images from dopamine axon terminals (anti-TH; green) with (bottom) or without (top) overnight 1uM sulpiride treatment; and their respective glycogen level content (color-code from dark violet = low levels to yellow = high levels; same scale top and bottom). (B) Average glycogen expression from axonal terminals normalized to control (gray dots; n = 25 neurons from 3 cultures), shows that glycogen storage levels decrease by a 45.4% ± 5.9 when neurons were treated with sulpiride (green dots; n = 21 from same cultures than control). (C) Average traces at the second round of stimulation (50AP, 10Hz) in absence of glucose in control neurons (gray trace; n = 29) versus neurons treated with sulpiride (green trace; n = 7). Responses are normalized to the peak. Vertical dashed line represents three times t measured at 5mM glucose response (data not shown). Horizontal dashed line represents 50% retrieval of the exocytic signal. (D) Number of rounds of stimulation before EB exceeds 50% is smaller in sulpiride treated neurons (green dots) than control neurons (gray dots): mean ± SEM: 1.3 ± 0.18 rounds versus 18.66 ± 1.69 rounds. Where 8 out 29 dopaminergic neurons never arrested for all 30 rounds of stimulation. (E) Average axonal ATP changes to 600 APs, 10Hz stimulation in neurons expressing iATPSnFR2.0 in the absence of glucose, normalized to pre-stimulation baseline. Control (gray trace, n = 14) vs sulpiride treated neurons (green, n = 6) shows that D2-autoreceptor signaling controls glycogen storage, and that sulpiride reduces this storage to levels insufficient for sustaining synaptic vesicle function during and after activity. (F) Comparison of ATP percentage before (Baseline), during (Drop) and after (Recovery) electrical activity in 5mM glucose (Gluc 5) and after 5min in a glucose free saline solution (Gluc 0). ^****^P<0.0001, ^***^P<0.001, ^**^P<0.01, non-significant as ns, Wilcoxon-Mann-Whitney test.

## Discussion

The basis for selective vulnerability of certain cell types in the brain remains a central question for understanding the molecular origins of neurodegeneration. This question is particularly relevant in PD, as the degree of degeneration of dopamine neurons in the Substantia Nigra pars compacta (SNc), are strongly correlated with duration of the disease (28, 29). Several important clues point to the possibility that PD, and the selective vulnerability of dopamine neurons may arise from bioenergetic deficits. Quantitative analysis of glucose metabolism suggests that brain hypoglycemia may be an early predictor of PD (30) and genetic studies in humans in the last three decades has successfully identified over twenty monogenic drivers of familial PD including many that are implicated either directly or indirectly in impacting cellular bioenergetics. Several of the disease driving genetic variants participate in mitochondrial quality control (31) e.g. Parkin (*PARK2*), and Pink1 (*PARK6*) (31) or are themselves *bona fide* mitochondrial proteins e.g. CHCHCD2 (*PARK22*) (32). Furthermore, recent evidence has accumulated that several other genetic drivers of PD either impair neuronal glycolysis e.g. DJ-1 (*PARK7*) and PGK1 (*PARK12*) or create a metabolic burden in axons (17). An FDA approved treatment of benign prostate hyperplasia, terazosin, was discovered to have an off-target enhancement of glycolysis (33). Terazosin in turn has proven remarkably effective in several animal models of PD (34) and retrospective epidemiology has shown that terazosin significantly reduces the risk of developing PD in humans compared to different BPH treatments with the same intended molecular target (35). These findings strongly support the idea that both monogenic and idiopathic PD have a strong bioenergetic component, in turn pointing to the need to examine the bioenergetic properties of mid-brain dopamine neurons. The data we show here demonstrate that *in vitro*, contrary to expectations, nerve terminal function in primary dopamine neurons is much more resilient to fuel withdrawal than primary hippocampal neurons. We traced the origin of this resilience to the fact that primary dopamine neurons under these conditions were reliant on stored glucose, in the form of glycogen, but that the availability of such a store was itself dependent on whether the neurons had previously activated their D2Rs. Previous work has shown that glycogen was protective against hypoxia in neurons (2) indicating that when available glycogen would be protective. Remarkably this resilience to fuel withdrawal comes at a price, as if either the glycogen stores are depleted (Fig. 4D) or access to them is blocked (Fig. S2C), primary dopamine neurons are more sensitive to fuel restriction. These data lead us to propose that the combination of a reliance on glycogen coupled with the autocrine nature of glycogen control may explain the general susceptibility of dopamine neurons in SNc to metabolic stressors, as once these neurons stop secreting sufficient dopamine such that resting levels in the striatum become depleted, this in turn leads to a depletion of glycogen stores resulting effectively to an positive feedback that leads to even greater metabolic stress in dopamine neurons, exacerbated nerve terminal function and further dopamine depletion.

In muscle, it has long been appreciated that glycogen stores serve as an important component of the “fight or flight” response, whereby adrenaline secretion drives glycogenolysis in turn boosting the bioenergetic capacity of the muscle. Our work demonstrates that neurons appear to also make use of neuromodulation to control their bioenergetic capacity by providing a signal to build energy reserves. At present we do not know if neurons, and in particular dopamine neurons, utilize neuromodulation (presumably by elevating cAMP) to facilitate glycogenolysis or if they simply rely on an intrinsic Ca^2+^modulation (perhaps via a Ca^2+^-sensitive adenylate cyclase) of this activation. Given the widespread expression of GPCRs in the brain, it is tempting to speculate that part of this may be used to locally gate both storage (via Gi activation) and use of energy reserves (via Gs activation), that in turn will help control cognitive function.

We do not understand at this point the molecular basis for the hypersensitivity of dopamine neurons to metabolic stress when deprived access to glycogen. It may reflect a fundamental difference some aspect of the molecular organization the glycolytic machinery that is well adapted to direct delivery of phosphorylated glucose from the glycogen store, but poorly adapted to extracellular glucose delivery. Further studies to understand this point are warranted as well as the general applicability of these findings to other neuron types both *in vitro* and *in vivo*. Our work is consistent with the observation that specific loss of glycogen machinery in neurons *in vivo* degrades neuronal function *in vivo* in rodents and worms (36, 37) suggesting that glycogen use by neurons is a fundamental property and means of regulating bioenergetic capacity in the brain.

## Materials and Methods

All protocols for this study have been deposited at: dx.doi.org/10.17504/protocols.io.ewov11q42vr2/v1

### Reagents

Chemical reagents were purchased from MilliporeSigma. Glycogen Phosphorylase Inhibitor (GPI) was purchased from Cayman Chamical (18578). Sulpiride was from MilliporeSigma (S7771). Antibodies used are anti (a)-TH in rabbit (Millipore, 657012, RRID: AB_2201407, 1:1000), a-TH in sheep (Millipore, AB1542, RRID: AB_90755, 1:1000), a-tubulin-βIII (R&D Systems, MAB1195, RRID: AB_357520, 1:1000), a-Glycogen Phosphorylase (Synaptic Systems, 255 003, RRID:AB_2619966, 1:500), a-Glycogen Synthase 1 (Proteintech, 10566-1-AP, RRID: AB_2116401, 1:500), IV58B6 a-Glycogen (provided by Dr. M. Gentry, 1:500), and a-TagFP nanobody conjugated with ATTO488 (NanoTag Biotechnologies, N0502-At488-L, RRID: AB_2744623; 1:500). Alexa Fluor–conjugated fluorescent secondary antibodies were obtained from Life Technologies and used at 1:500.

### Animals

All experiments involving animals were performed in accordance with protocols approved by the Weill Cornell Medicine Institutional Animal Care and Use Committee. Neurons were derived from Sprague-Dawley rats (Charles River Laboratories strain code: 001, RRID: RGD_734476) of either sex on postnatal days 0 to 1.

### Plasmids

The following previously published DNA constructs were used: syphy-pH (Addgene, plasmid # 220506, RRID: Addgene_220506). For this study, these new plasmids were generated: PYGB-KD (referred here as GP-KD, targeting sequence CCTGTATCCCAATGACAATTT, Addgene, plasmid # 242946, RRID: Addgene_242946), THP Synaptophysin-pH (Addgene, plasmid # 242903, RRID: Addgene_242903) and THP cyto-iATPSnFR2-miRFP670nano3 (Addgene, plasmid # 242904, RRID: Addgene_242904).

### Primary neuronal culture

Ventral midbrain primary neuronal culture and transfections were applied as described (38). Hippocampal primary neuronal culture and transfections were applied as described (39).

### Live-cell imaging

Live-cell imaging was performed as described (40). Neurons were continuously perfused at 0.1 ml/min with a Tyrode’s solution 119 mM NaCl, 2.5 mM KCl, 5 mM glucose, 50 mM HEPES, 2mM CaCl2, 2 mM MgCl2, 50 μM DL-2-amino-5 phospho-novaleric acid (AP5), and 10 μM 6-cyano-7-nitroquinoxaline-2,3-dione (CNQX), adjusted to pH 7.4. For free glucose experiments, Tyrode’s solution osmolarity was adjusted with 55mM HEPES. For sulpiride experiments, cultured dishes were incubated between 12 to 16 hours with final drug concentration of 1uM. When GPI was used, it was added at 10uM final concentration.

### Axonal ATP Image Analysis

iATPSnFR2 image processing and analysis was performed as described (41). Time series of imaging pairs (miRFP670nano3 and iATPSnFR2) were split into two independent image series using a custom-written Fiji routine to facilitate analysis. ATP signals are reported as a ratio between iATPSnFR2: miRFP670nano3. Images were analyzed using the ImageJ plug-in Time Series Analyzer V3 where 40 to 60 circular ROIs of radius 1 μm corresponding to axonal terminals expressing the iATPSnFR2 were selected in the miRFP670nano3 channels, blind to the iATPSnFR2 channel, and then background subtracted. Image loading and posterior raw data saving were automatized using a homemade Python code for Fiji. Average ROIs signals were analyzed using homemade script routines in Igor-pro v6.3.7.2 (WaveMetrics, Lake Oswego, OR, USA). ATP ratio signal was calculated by averaging all individual ROIs per neuron and normalized to the baseline.

### pHluorin measurements

pHluorin image processing and analysis was performed as described (42). For pHluorin experiments, nerve terminals responding to the first AP train in 5mM glucose were selected and background subtracted. In case of the repeated stimulation assay, the traces are reported as percentage of total sensor expression by subtracting the initial fluorescence signal before stimulation and normalized to total sensor fluorescence, revealed by the 50 mM NH_4_Cl Tyrode’s solution. For single-train experiments, the traces are normalized to the fluorescence peak during the AP train.

### Immuno-staining

Immunofluorescence staining was performed as described (43). Neurons were fixed with 4% paraformaldehyde, quench PFA with 50mM NH_4_Cl solution, permeabilized with 0.25% Triton X-100, and blocked for 10 min at room temperature with 5% BSA. Primary antibodies were diluted with 5% BSA and incubated with the cells at room temperature for 1 hour. After 3× 5-min washes in phosphate-buffered saline (PBS), cells where incubated with secondary antibodies, followed by additional 3× 5-min washes in PBS. Immunofluorescence images were taken by the camera in a similar way as live-cell imaging.

### Image analysis and statistics

Images were analyzed using the ImageJ plug-in Time Series Analyzer V3 where 20 to 30 circular regions of interest (ROIs) of radius ∼1 um corresponding to synaptic boutons expressing the pHluorin (as determined with NH4Cl perfusion) or iATPSnFR2 (with miRFP670nano3 positive) were selected, and the fluorescence was measured over time. Image loading and posterior raw data saving were automatized using a homemade Python code for Fiji (RRID: SCR_002285). ROIs signals were analyzed using homemade script routines in Igor-pro (Wavemetric, RRID: SCR_000325). Results of group data analysis are presented as mean ± SEM. When analyzing means, *P* values are based on the nonparametric Wilcoxon-Mann-Whitney test. *P* < 0.05 was considered significant and denoted with a single asterisk, whereas *P* < 0.01, *P* < 0.001, and *P* < 0.0001 are denoted with two, three, and four asterisks, respectively. The *n* value, indicated in the figure legends for each experiment represents the number of cells imaged, otherwise indicated.

## Acknowledgments

We thank members of the Ryan laboratory for their valuable suggestions and input on this work. We thank Drs. Otto Baba (Tokushima University) and Ben J. Appelmelk (Amsterdam University Medical Center) for providing the hybridoma cell line producing the anti-glycogen monoclonal antibody. This research was funded in part by NIH to NS11739 (T.A.R.) and R35NS116824 (M.S.G.) and in part by Aligning Science Across Parkinson’s (ASAP-000580, ASAP-024404) through the Michael J. Fox Foundation for Parkinson’s Research (MJFF).

## Data and materials availability

The data, code, protocols, and key lab materials used and generated in this study are listed in a Key Resource Table alongside their persistent identifiers at 10.5281/zenodo.16877070. All data cleaning, preprocessing, analysis, and visualization was performed using Fiji (RRID:SCR_002285) and IGOR-Pro (WaveMetrics, RRID: SCR_000325).

## Supporting Information for

**Fig. S1.**
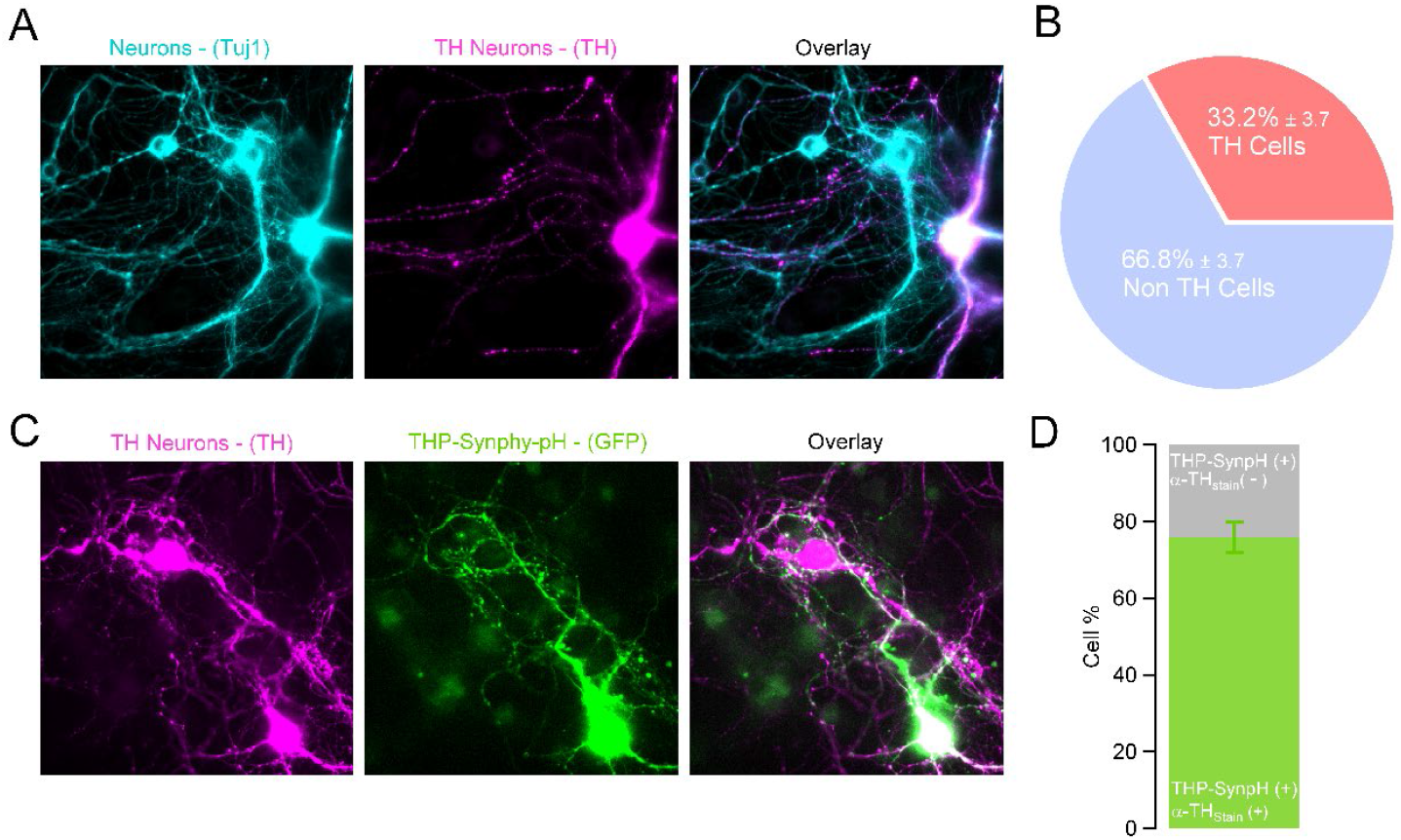
Dopaminergic neurons comprise about one-third of the ventral midbrain culture and can be selectively targeted via the TH promoter. (A) Representative image of cultured neurons (anti-Tuj1; pseudo-colored cyan) that are identified as dopaminergic (TH+) neurons (anti-TH; pseudo-colored magenta). Overlay in right panel. (B) Approximately, one-third of cultured neurons are dopaminergic (n = 11 dishes). (C) Representative field of view showing that one of the dopaminergic neurons (anti-TH; pseudo-colored magenta) is transfected with Syphy-pH under TH promoter (anti-GPF; pseudo-colored green, publication). Overlay in right panel. (D) TH 75.8 ± 4% of neurons transfected with the TH promotor subsequently were identified as dopaminergic (n = 21 dishes).

**Fig. S2.**
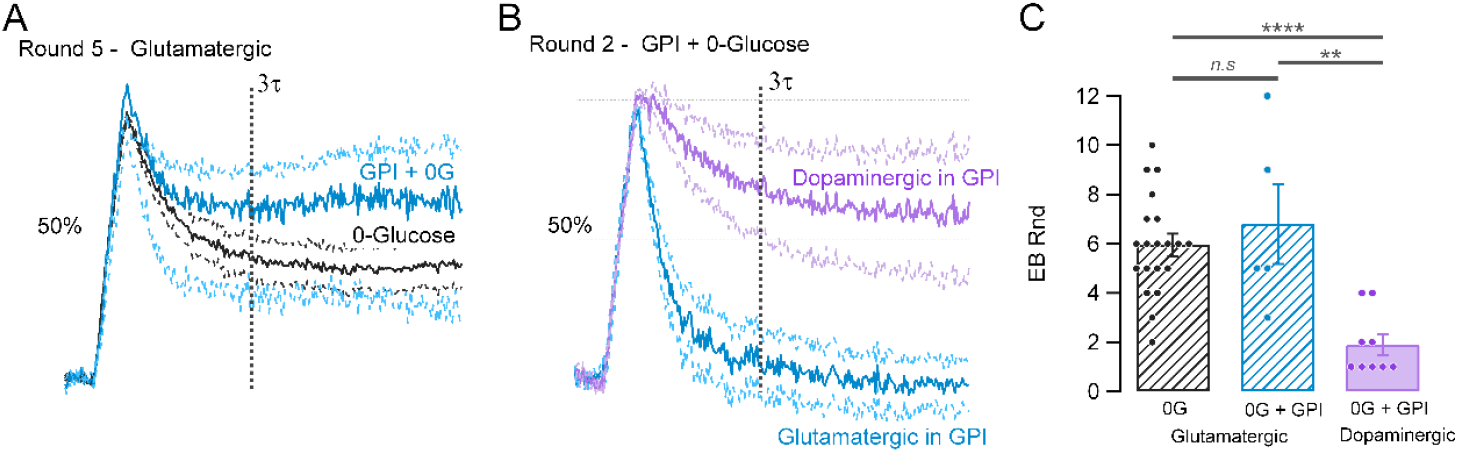
Glutamatergic neurons do not relay on glycogen under hypoglycemic conditions. (A) Average traces at the fifth round of stimulation (50AP, 10Hz) in absence of glucose in control glutamatergic neurons (gray trace; n = 20) versus glutamatergic neurons treated with GPI (blue trace; n = 5) shows that both populations are equally functionally impaired. (B) Second round of stimulation comparison between dopaminergic versus glutamatergic neurons treated with GPI shows that early on dopaminergic SV recycling is affected due to lack of access to glycogen, whereas glutamatergic neurons are not affected by GPI treatment. (A-B) Responses are normalized to the peak. Vertical dashed line represents three times t measured at 5mM glucose response. Horizontal dashed line represents 50% retrieval of the exocytic signal. (C) Number of rounds of stimulation before EB exceeds 50% is equally higher in glutamatergic neurons with or without GPI than in Dopaminergic neurons in GPI.

**Fig. S3.**
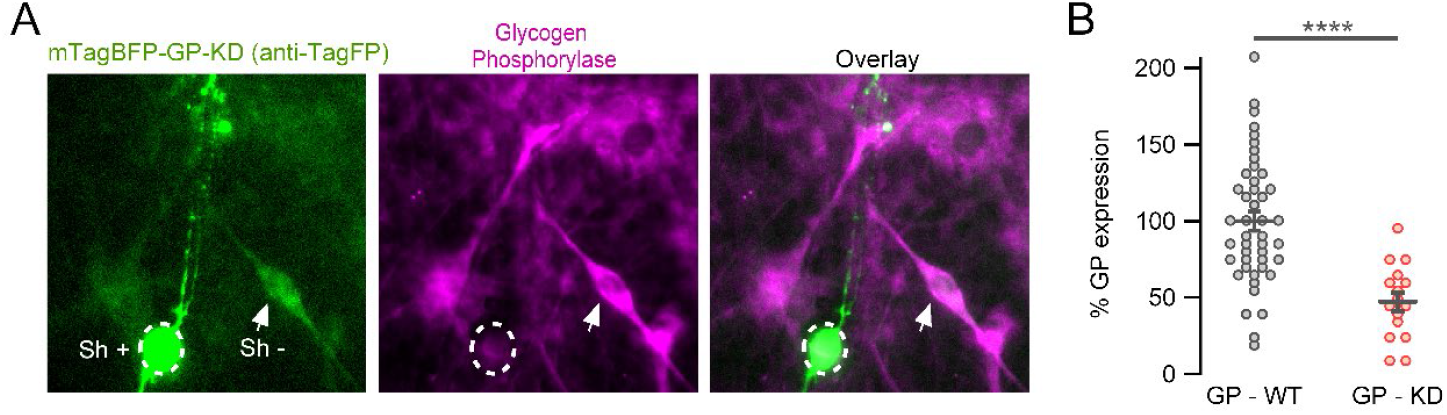
Expression of a ShRNA targeting Glycogen phosphorylase resulted in ∼35% suppression of GP expression. (A) Representative image from a neuron expressing an mTagBFP-GP-Sh (stained with FlutoTag-X2 anti-TagFP Atto488). BFP-positive cell body (dashed white circle) show much lower GP immunofluorescence when compared with neighboring non-transfected cell body (white arrow). (B) Average expression of GP in cell bodies transfected with mTagBFP-GP-Sh (n = 16) normalized to their respective neighboring non-transfected cell bodies (n = 45) from 3 cultures. Respectively as: mean ± SEM: 47.25%± 6.0 vs 100% ± 6.2. ^****^P<0.0001, Wilcoxon-Mann-Whitney test.

**Fig. S4.**
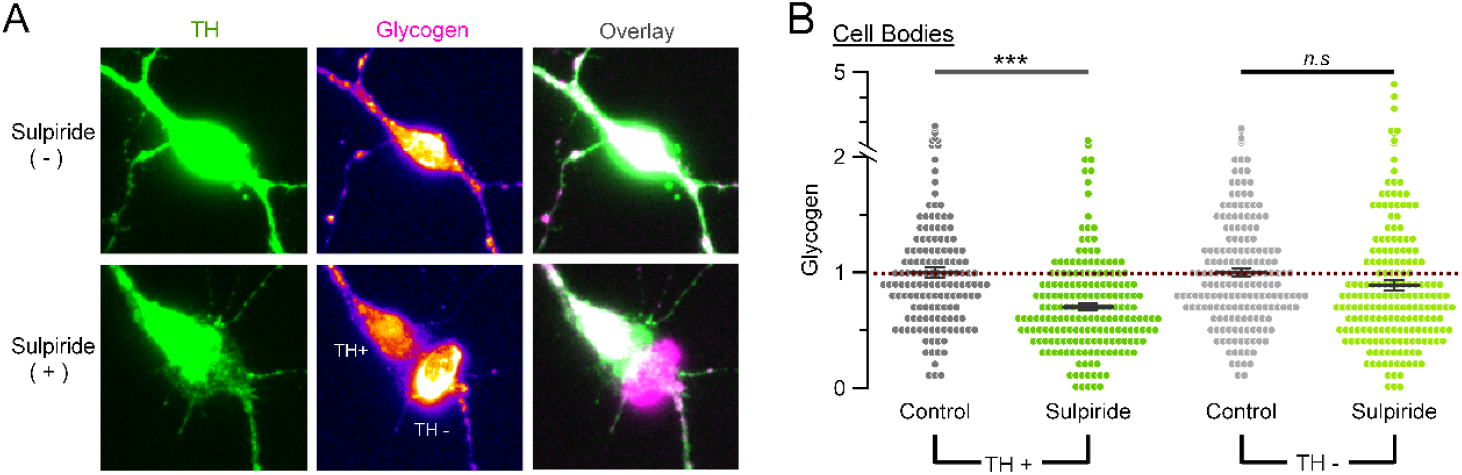
Sulpiride decrease glycogen levels exclusively in dopamine neurons, which are modulated by D2 auto-receptors. (A) Representative images of cell bodies from dopamine neurons (anti-TH, green) with (bottom) or without (top) overnight 1uM sulpiride treatment; and their respective glycogen level content (color-code from dark violet = low levels to yellow = high levels; same scale top and bottom). The glycogen field of view image treated with sulpiride shows a comparison between a TH+ versus a non-TH neuron. (B) Average glycogen expression from cell bodies normalized to control of TH+ (dark gray dots; n = 146) and to control of TH-populations (light gray dots; n = 183), shows that glycogen storage levels decrease by a 29.76% ± 2.94 only in neurons that are dopaminergic (TH+) and that were treated with sulpiride (dark green dots; n = 193 from same cultures than TH+ control). ^***^P<0.001, non-significant as ns, Wilcoxon-Mann-Whitney test.

